# Whole genome sequence of Mapuche-Huilliche Native Americans

**DOI:** 10.1101/252619

**Authors:** Elena A. Vidal, Tomás C. Moyano, Bernabé I. Bustos, Eduardo Pérez-Palma, Carol Moraga, Alejandro Montecinos, Lorena Azócar, Daniela C. Soto, Eleodoro Riveras, Mabel Vidal, Alex Di Genova, Klaus Puschel, Peter Nürnberg, Stephan Buch, Jochen Hampe, Miguel L. Allende, Verónica Cambiazo, Mauricio González, Christian Hodar, Martín Montecino, Claudia Muñoz-Espinoza, Ariel Orellana, Angélica Reyes-Jara, Dante Travisany, Paula Vizoso, Mauricio Moraga, Susana Eyheramendy, Alejandro Maass, Giancarlo V. De Ferrari, Juan Francisco Miquel, Rodrigo A. Gutiérrez

## Abstract

**Background:** Whole human genome sequencing initiatives provide a compendium of genetic variants that help us understand population history and the basis of genetic diseases. Current data mostly focuses on Old World populations and information on the genomic structure of Native Americans, especially those from the Southern Cone is scant.

**Results:** Here we present a high-quality complete genome sequence of 11 Mapuche-Huilliche individuals (HUI) from Southern Chile (85% genomic and 98% exonic coverage at > 30X), with 96–97% high confidence calls. We found approximately 3.1×10^6^ single nucleotide variants (SNVs) per individual and identified 403,383 (6.9%) of novel SNVs that are not included in current sequencing databases. Analyses of large-scale genomic events detected 680 copy number variants (CNVs) and 4,514 structural variants (SVs), including 398 and 1,910 novel events, respectively. Global ancestry composition of HUI genomes revealed that the cohort represents a marginally admixed population from the Southern Cone, whose genetic component is derived from early Native American ancestors. In addition, we found that HUI genomes display highly divergent and novel variants with potential functional impact that converge in ontological categories essential in cell metabolic processes.

**Conclusions:** Mapuche-Huilliche genomes contain a unique set of small– and large-scale genomic variants in functionally linked genes, which may contribute to susceptibility for the development of common complex diseases or traits in admixed Latinos and Native American populations. Our data represents an ancestral reference panel for population-based studies in Native and admixed Latin American populations.

## Background

Sequencing complete human genomes has greatly expanded the knowledge of our genetic diversity, providing insights into the evolutionary history of man and the bases of human diseases. Large-scale genomic initiatives such as HapMap [1], the 1000 Genomes Project (1kGP) [2] or the ExAC initiave [3] have revealed that individuals from multiple populations carry different profiles of rare and common variants that differ substantially among human continental groups. While current high-coverage full genome efforts have mostly focused on Old World populations (Europeans, Asians and Africans), there is still limited information concerning the genetic structure of Native American groups [4].

Genome-wide sequence data from ancient and present-day humans has been described from Greenland, Arctic Canada, Alaska, Aleutian Islands and Siberia and used to understand migration pulses into the Arctic regions of America [5, 6]. Likewise, whole genome/exome and large-scale genotyping data has been used to study the genetic history, multiple streams of migration and population-genomic variables that underlie patterns of deleterious variation in African, Asian, European, and Native American ancestry in populations of Latin America and the Caribbean [7–10], as well as the Pacific Northwest [11]. More recently, studies on the demographic history and population structure of admixed South American Latinos has been reported with the aid of genome-wide genotyping technologies [12–14]. Nevertheless, whole genome sequencing efforts aimed to identify genetic variants that may influence susceptibility to develop complex common disorders affecting modern Native American and admixed American populations are still scarce.

Mapuche-Huilliches descend from early hunter-gatherers who colonized the subcontinent about 15000 years ago and are the modern representatives of one of the most prominent indigenous groups in the Southern Cone of South America [15]. Here we sequenced at high-coverage and analyzed the complete genome of 11 individuals belonging to a native Mapuche-Huilliche population from Southern Chile. Addressing genetic contributions from Native populations represents a major step towards understanding the genetic basis for common traits or disorders in current admixed populations.

## Results

### Overview of Mapuche-Huilliche genomes

Individuals sequenced belong to a community living in Ranco Lake (latitude 40°13’27.62”S, longitude 72°22’50.16”W), in the Los Rios region of Southern Chile (**Additional file 1: Figure S1**). A common difficulty encountered while studying Native American genetic history is the admixture with individuals from Europe and Africa that occurred since the arrival of Europeans to America in 1492. For example, Spanish conquerors arrived in Chile in the sixteenth century and began interbreeding with native females, primarily of the Mapuche-Huilliche group, giving birth to the current Chilean population [12, 16, 17]. Therefore, to ensure *a priori* as little admixture as possible, 11 individuals from the Mapuche-Huilliche community (10 females and 1 male) with > 3 surnames of Mapuche origin, ABO type O and Rh+, the most common blood type in southern Cone Native Americans, and that were homozygous for a SNV located 13.9 kb upstream of the human lactase gene (*LCT*: C>T-13910; rs4988235), which determines the lactase nonpersistent state in Native Americans [18], were selected for sequencing (Table 1), hereinafter referred as “HUI” cohort. HUI DNA samples were sequenced using the combinatorial probe-anchor ligation and DNA nanoarray technology of Complete Genomics [19]. We obtained an average of 85% genomic and 98% exonic coverage of at least 30X, with 97% and 98% high confidence calls, respectively.

**Table 1.** Details of HUI individuals selected for this study and genome sequencing statistics.

| **Assembly ID** | **Sex** | **Mapuche surnames** | **Age at screening** | **Called genome fraction** | **Genome coverage 30X** | **Exome coverage 30X** | **Mapping yield (Gb)** |
| --- | --- | --- | --- | --- | --- | --- | --- |
| GS000011194 | F | 4 | 22 | 0.97 | 0.87 | 0.98 | 180.69 |
| GS000011195 | F | 4 | 47 | 0.97 | 0.87 | 0.98 | 181.74 |
| GS000011196 | F | 4 | 42 | 0.97 | 0.85 | 0.98 | 180.88 |
| GS000011198 | F | 4 | 46 | 0.97 | 0.84 | 0.98 | 180.61 |
| GS000011200 | F | 4 | 18 | 0.97 | 0.84 | 0.98 | 177.51 |
| GS000011201 | F | 4 | 53 | 0.96 | 0.83 | 0.98 | 175.46 |
| GS000011215 | F | 4 | 43 | 0.97 | 0.88 | 0.98 | 180.44 |
| GS000012210* | M | 4 | 49 | 0.96 | 0.86 | 0.99 | 266.52 |
| GS000012242* | F | 4 | 27 | 0.96 | 0.92 | 0.99 | 357.46 |
| GS000020403 | F | 4 | 39 | 0.97 | 0.82 | 0.98 | 151.00 |
| GS000020711 | F | 3 | 60 | 0.97 | 0.84 | 0.98 | 177.72 |

Called genome fraction: Fraction of the reference genome with full (diploid) calls in the sequenced sample following assembly; Genome coverage 30X: Fraction of the reference genome bases where coverage is greater than or equal to 30X; Exome coverage 30X: Fraction of the reference exome bases where coverage is greater than or equal to 30X; Mapping yield: Total base-pairs of sequence reads mapped to the reference genome. Samples marked with an asterisk were sequenced twice, thus they have greater mapping yields and average coverage than the other samples.

**Table 2.**
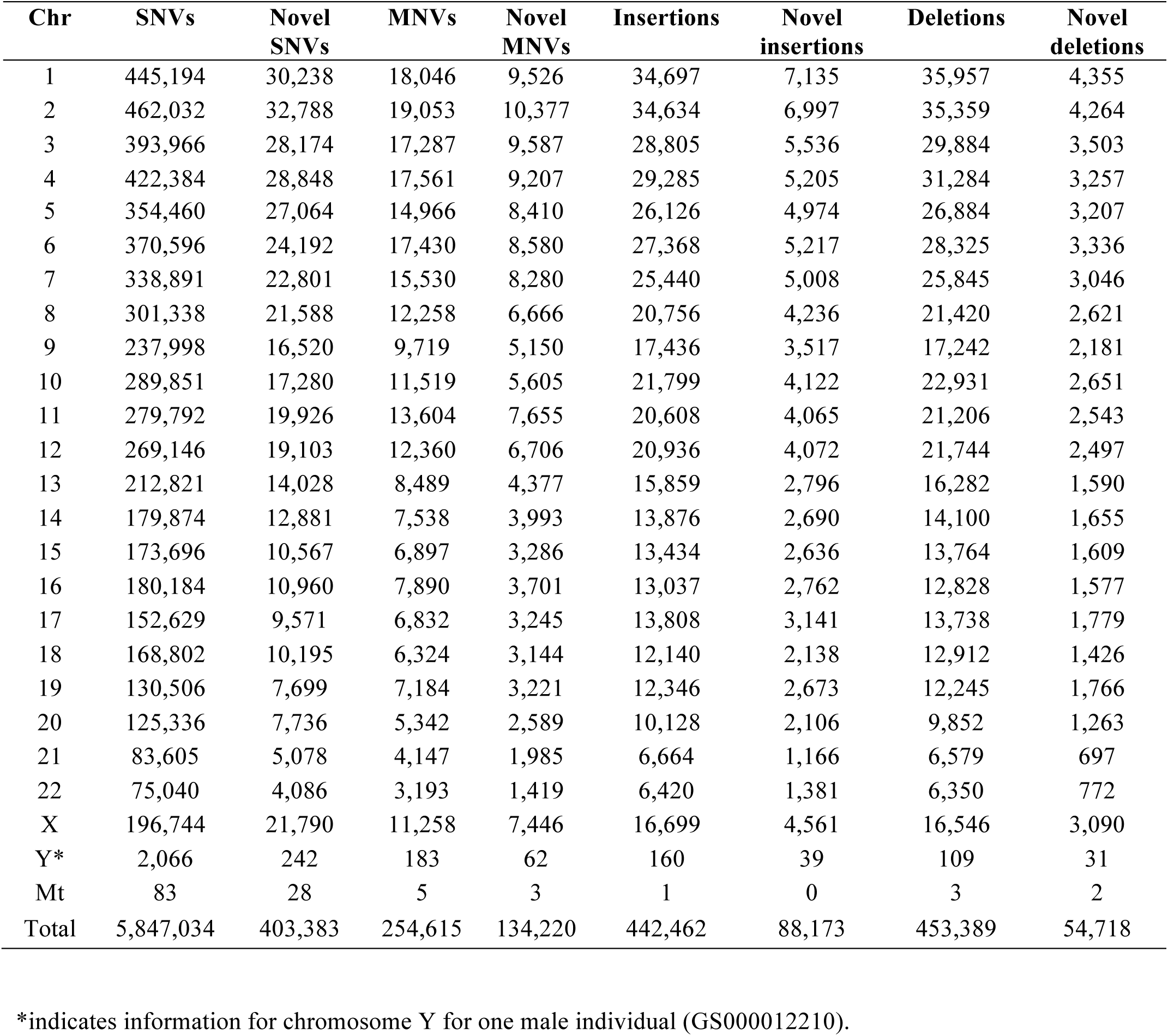
Total variants identified in HUI individuals listed by chromosome.

Approximately 3.1×10^6^ single nucleotide variants (SNVs) were determined for each individual, with a total of 5,847,034 SNVs in the cohort (**Additional file 1: Table S1**). We found a high level of concordance (99.70%) in the SNV calling rate between genome sequencing and genotyping results obtained using the Illumina Infinium Human Core Exome BeadChip on the same individuals (**Additional file 1: Table S2**). Since the Illumina chip is focused only in exonic and gene-surrounding variants, we determined the genome-wide transition versus transversion mutations (Ts/Tv) ratio for all variants to assess for the presence of false positive calls. We observed that the Ts/Tv ratio in sequenced genomes is 2.1:1 (**Additional file 1: Table S3**), in agreement with the 1kGP expected ratio of 2:1 [2], further confirming the good quality of variants. Identity by descent (**Additional file 1: Table S4**) and inbreeding (**Additional file 1: Table S5**) analyses indicated that individuals sequenced were not closely related or inbred. A genome-wide summary of main genetic elements in HUI individuals is presented in Fig. 1.

**Fig. 1.**
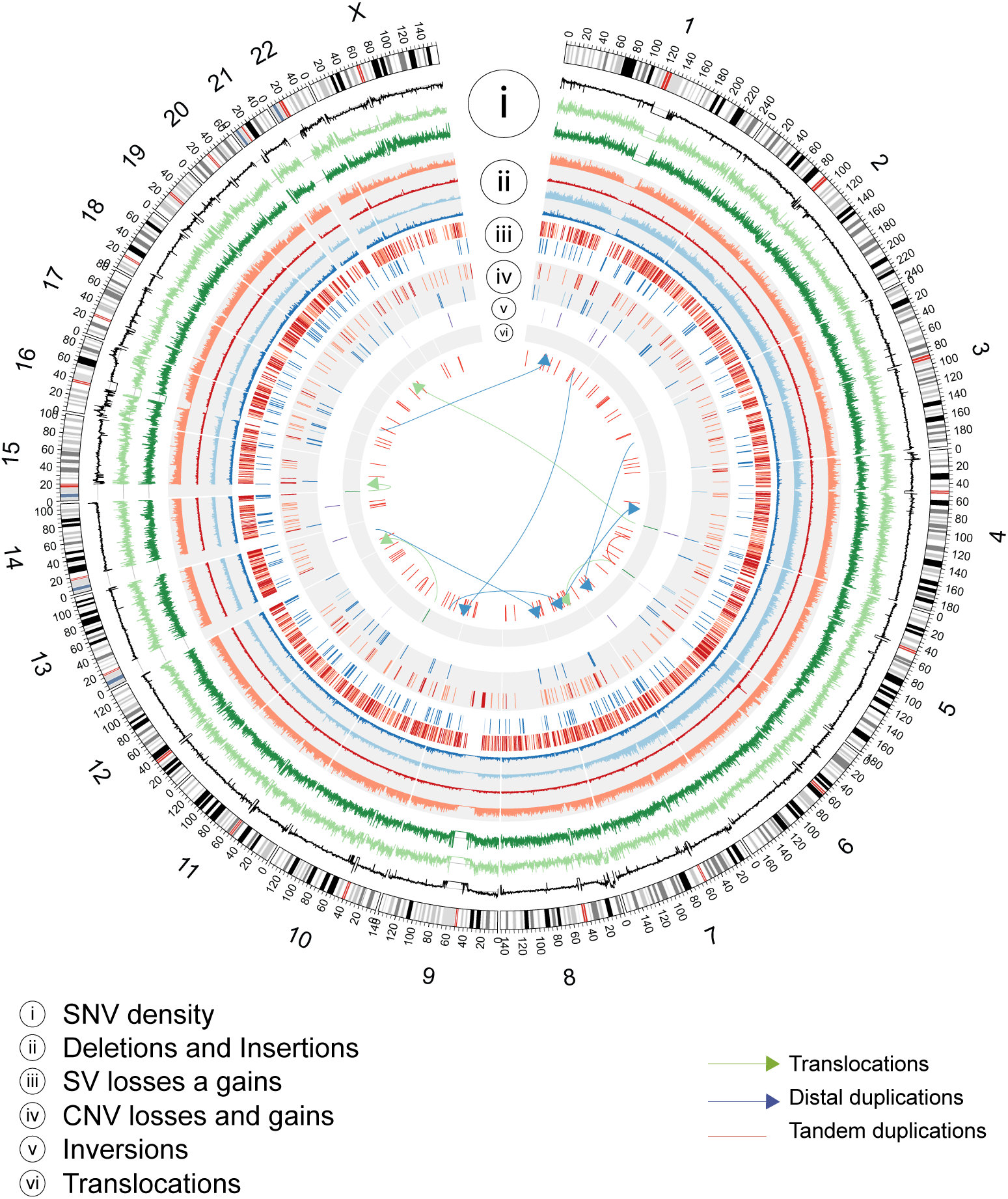
Genetic and structural variants in Mapuche-Huilliche genomes. Circos plot of the spatial distribution of SNV densities (i), deletions and insertions (ii), structural variant (SV) loses and gains (iii), copy number variant (CNV) losses and gains (iv), inversions (v) and translocations (vi). Light or dark colors in different tracks indicate known or novel variants, respectively. Tandem (red lines) and distal duplications (blue arrows) are shown within the inner circle of the plot. Translocation events are shown as green arrows.

We identified 403,383 (6.9%) novel SNVs that are not included in dbSNP build 144 release or do not have a reported frequency either in the 1kGP-phase 3 database [2], the Exome Sequencing Project [20] or the Exome Aggregation Consortium (ExAc) [21] (**Additional file 1: Table S1**). To reduce the amount of variants with missing calls, we applied a calling rate (CR) threshold of 90% giving a total number of 321,803 SNVs, including an excess of individual privative variants (1/22 alleles; 256,550 SNVs, 79.7 %; **Additional file 1: Fig. S2a**), in agreement with the current literature [22]. A number of 175,897 novel SNVs fell in intergenic regions (54.66%), 118,777 in introns (36.91%) and 1,769 in the coding portion of the genome (0.55%, exonic) (**Additional file 1: Fig. S2b**). Likewise, we observed 88,173 (19.92%) insertions and 54,718 (12.06%) deletions that are novel and observed in at least 1 individual. In addition, analyses of large-scale genomic events detected 680 copy number variants (CNVs) and 4,514 structural variants (SVs), including 398 and 1,910 novel events (**Additional file 1: Table S6 and S7**, respectively), that did not overlap any region reported in the CNV map from the Database of Genomic Variants [23] or the 1kGP-phase 3 release [24]. We found 1,096 genes partially or completely overlapped by ≥1 CNVs or SVs in at least one HUI sample (**Additional file 1: Fig. S3a** and **Additional file 2: Table S8**) and 37 genes consistently affected by novel events in all HUI genomes analyzed. Interestingly, some of these large structural events alter the coding sequence of genes and thus may have a potential functional impact (**Additional file 1: Fig. S3b**).

### Global ancestry composition of HUI and Chilean Latino genomes

We used ADMIXTURE software [25] to determine ancestry composition of HUI genomes by comparing a set of 105,252 SNVs shared with those present in the complete set of samples of the 1kGP-phase 3 (n = 2,504 individuals) [2]. In addition, we included a panel of 1,191 Chilean Latinos genotyped by the Affymetrix Axiom World Array LAT 1, to represent the general Chilean population. We ran ADMIXTURE from K = 1 to K = 15 models and obtained near minimum values beyond K = 5 (CV error = 0.51976, **Additional file 1: Fig. S4a**) and a minimum cross-validation error at K = 10 (CV error = 0.51868, **Additional file 1: Fig. S4b**). First, we explored a continental model that considers 5 ancestry components (K = 5), which includes the super-populations from European (EUR), East Asian (EAS), South Asian (SAS), African (AFR) and admixed American (AMR) ancestries from 1kGP-phase 3 data. We found that HUI samples have a high degree of admixed American ancestry (average = 93.8%) with a minimal contribution of other founder populations (Fig. 2a, Top), which validates our ascertainment scheme for selection of individuals to be sequenced. In agreement with recent data regarding the ancestry and complex demographic history of South America [13], the Chilean Latino panel behaves as an admixed population with a strong genetic contribution of both EUR and AMR ancestries (average = 49.97% and 45.54%, respectively). Other AMR cohorts from 1kGP-phase 3, such as Peruvians from Lima (PEL), Mexicans from Los Angeles (MXL), Colombians from Medellin (CLM) and Puerto Ricans in Puerto Rico (PUR), showed important contributions from EUR and AMR and to a lesser extent from AFR and other super-populations. Second, when we ran ADMIXTURE considering the minimum cross-validation error (K = 10) we observed that all super-populations split into two major components within each specific cluster, as described before for K = 5 (Fig. 2a, Bottom). Notably, a large component of the AMR ancestry in PEL, MXL, CLM and PUR populations (dark gray) is not present in HUI genomes (average = 0.5%) and is marginally represented in Chilean Latino individuals (average = 6.9%, compared with 76.2% in PEL, 42.9% in MXL, 25.6% in CLM and 13.5% in PUR samples). These results suggest that HUI individuals and the broader Chilean cohort derive this genetic component from shared Native American ancestors with low genetic representation in other admixed American populations.

**Fig. 2.**
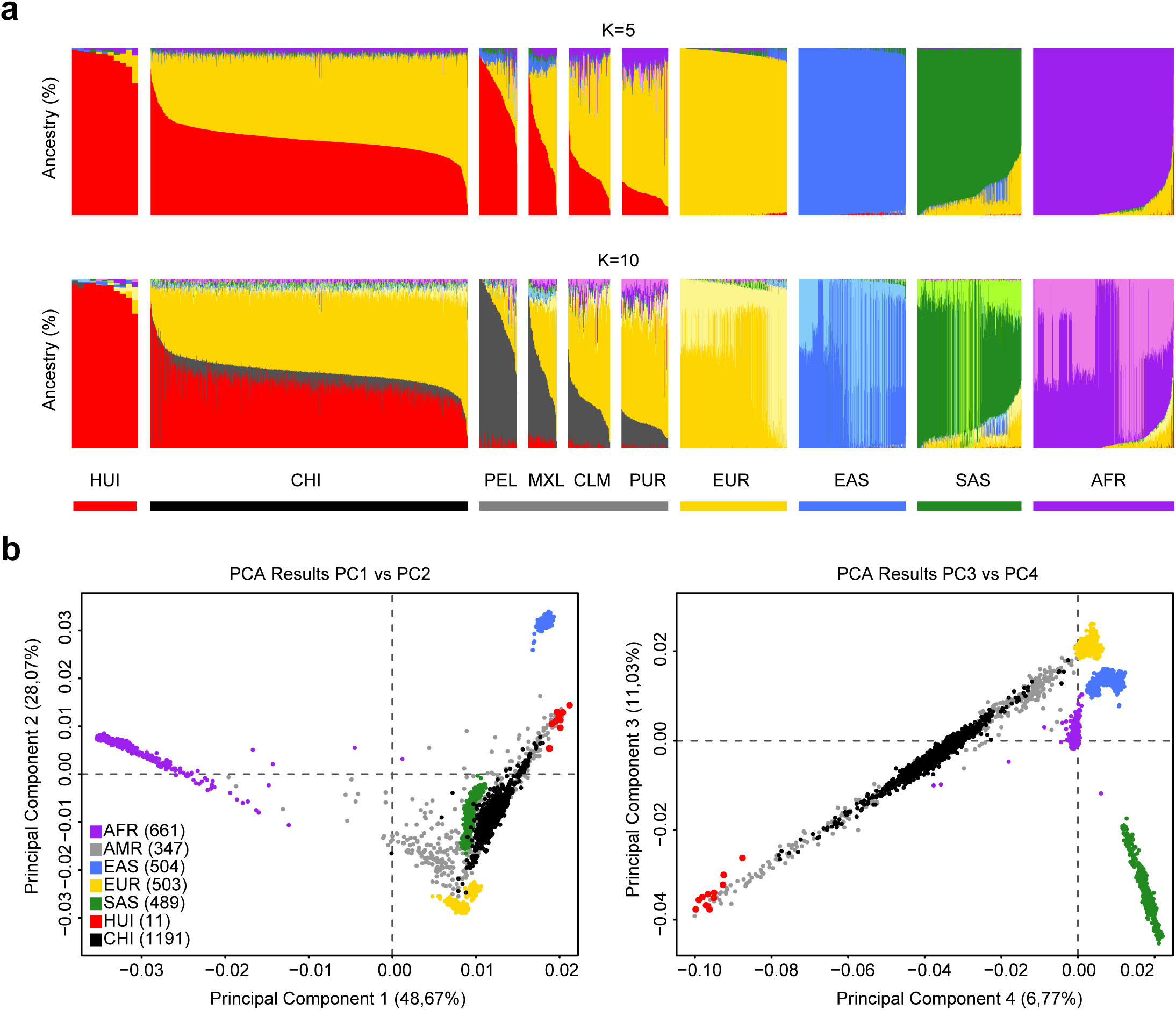
Ancestry analysis of HUI and Chilean Latino individuals. (**a**) ADMIXTURE plots for K = 5 (Continental model) and K = 10 (minimum error model). All 3,706 samples included are depicted as vertical thin bars colored by their corresponding ancestry percentage. HUI genomes are highlighted at the left with thicker bars followed by Chilean Latino genotyped individuals and samples included in 1kGP-phase 3, which are clustered in 5 super-populations (AMR, EUR, EAS, SAS and AFR). For K = 5, the colors were defined as follows: Red for “Amerindian”, yellow for “European”, blue for “East Asian”, green for “South Asian” and purple for “African”. For K = 10, light colors are used to show subcomponents within super-populations EUR, EAS, SAS and AFR. Grey color is used to represent the AMR component common to PEL, MXL, CLM and PUR populations but almost absent in HUI. Bottom thick bars define key colors used in the PCA. (**b**) Principal Component (PC) analysis including the same set of samples (colored dots) and markers. Color legend and number of samples belonging to each super population defined in (**a**) is provided in the legend inside brackets. Left Panel: PC1 vs. PC2,right panel: PC3 vs. PC4. Percentage of variance explained by each component is given in parenthesis in the corresponding axis.

To further explore the genetic structure of the HUI cohort, we used the same set of samples and SNVs and run a principal component analysis (PCA) using EIGENSTRAT [26] (Fig. 2b). PCA results clearly defined world population structure, showing clusters composed of African (AFR), European (EUR) and Asian (EAS and SAS) populations, while the admixed American superpopulation (AMR) was widely distributed between EUR and HUI individuals. Consistent with absence of significant recent admixture, our HUI cohort cluster in the PCA plot at one edge of the AMR samples (red vs. gray dots, respectively). As expected, the largest genetic distance existed between the AFR population and the rest of the groups. In turn, we observed that genotyped Chilean Latino samples (black dots) spread in a relatively narrow cluster that begins at the edge of HUI individuals and ends with individuals from the EUR founder population and overlapped most of the AMR cluster. However, in contrast to admixed Chilean Latinos, the AMR super populations were widely distributed between EUR, AFR and HUI populations. These results are in agreement with the admixture analysis showing that the Chilean Latino population exhibits minimal genetic contribution of other population beyond EUR and HUI/AMR, in accordance to the Chilean demographic history. Unlike other AMR populations with considerable contribution or African/Asiatic immigration (i.e. PUR; average = 15.0% of AFR contribution) [7, 27, 28], African and Asian ancestry in Chilean samples was almost negligible (average = 1.5% and 1.1%, AFR and EAS + SAS, respectively).

Analysis of mitochondrial DNA showed that all HUI individuals belong to the Native American haplogroups C and D, two of the major pan-continental founder haplogroups. The majority of genomes sequenced (7 out of 11) belong to the C1b haplogroup and 6 of them were assigned to the clade C1b13 (**Additional file 1: Fig. S5a**), which is a branch found mainly in the Southern Cone of South America between 38° and 42°S [15, 29]. While the other 4 individuals belong to the D haplogroup, 3 of them are in the D1g clade, which is found almost exclusively in the central-southern part of Chile and Argentina, and only one is in the D4h3a clade (**Additional file 1: Fig. S5b**), found mainly in the Southern Patagonia [15, 29]. These results are in agreement with the admixture data (K = 10, as described before) showing that the genetic component of the HUI cohort differs from the genetic component of other Native American populations living in the northern region of South America.

### Highly divergent and novel variants with potential functional impact in HUI genomes

Genetic variation explaining differential susceptibility to diseases or metabolic conditions derives mostly from studies in populations from developed countries, and more recently from genotyping admixed Latin American populations [12, 14, 30]. To identify highly divergent SNVs in HUI individuals compared to other world populations, as a measure of differentiation due to population structure, we determined pairwise fixation index (Fst) statistics [31] between HUI and 1kGP-phase 3 populations from Africa (AFR: YRI, LWK, GWD, MSL, ESN, ASW and ACB), Europe (EUR: CEU, IBS, GBR, FIN, TSI), America (AMR: CLM, PUR, MXL and PEL), Southern Asia (SAS: PJL, GIH, BEB, STU and ITU) and East Asia (EAS: CHB, JPT, CHS, CDX and KHV) [2]. At the population level, weighted Fst statistics revealed that HUI individuals are genetically closer to admixed American individuals (i.e. PEL, MXL, CLM and PUR), then with individuals from Eastern and Southern Asian and finally with European and African populations (Fig. 3a), in agreement with the settlement and population history of ancient Native Americans [7, 10]. Likewise, and at the variant level, Fst analyses allowed the identification of an average number of 165,746 SNVs surpassing the 95^th^ percentile of the genome-wide Fst distribution, which was used as a threshold of high divergence (HUI_95th_: average Fst cutoff of 0.51, vertical lines in Fig. 3b and **Additional file 1: Table S9**). The union of all Fst variants yielded a total number of 842,780 HUI_95th_ divergent SNVs, representing the whole genetic variability of HUI compared to other 1kG-P3 populations.

**Fig. 3.**
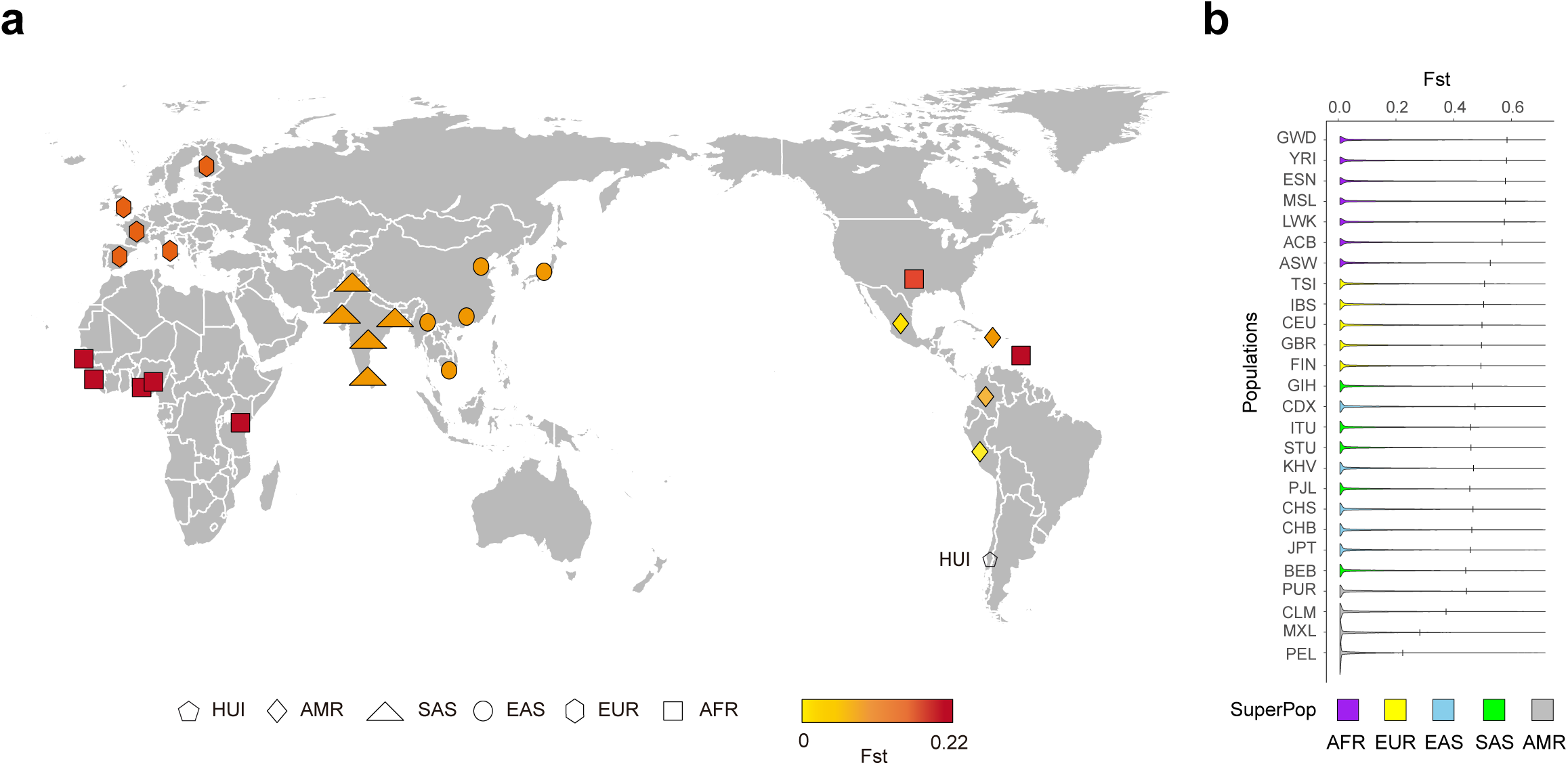
Highly divergent SNVs in HUI genomes. (**a**) World map showing all 26 populations from 1kGP-phase 3 coming from the 5 super populations (AFR, SAS, EAS, EUR and AMR) and their Weir and Cockerham’s Fst statistic (weighted Fst) from yellow to red according to their lower or higher divergence obtained from the comparison with the HUI sequenced individuals. (**b**) Violin plots comparing SNV density between HUI and other 26 populations from 1kGP-phase 3. Fst distributions are sorted by decreasing genetic divergence from HUI (top to bottom). Vertical bars on each population plot indicate 95th percentile cutoff used to identify highly divergent SNVs. SuperPop = Super populations from 1kGP-phase 3: AFR = Africans, AMR = Admixed Americans, ASN = Asians, EUR= Europeans.

To identify HUI divergent variants with potential functional impact (VPFIs) we followed a two-step variant/gene prioritization procedure, focused solely on exonic/exonic-splicing variants that may result in nucleotide changes affecting protein folding, structure or stability. First, Fst-derived HUI_95th_ variants detected in 2 or more populations were queried for functional impact using ANNOVAR [32] with the Combined Annotation Dependent Depletion (CADD) database [33], which quantitatively prioritize functional, deleterious and disease causal variants. We detected a total number of 529 intolerant HUI VPFIs annotated to 500 genes (**Additional file 3: Table S10**). Second, we searched for significant protein-protein interaction connectivity among these HUI VPFIs containing genes using the online database resource Search Tool for the Retrieval of Interacting Genes (STRING) [34], using only highest confidence interaction scores (0.9; probability that a predicted link exists between two proteins), and recovered 115 gene products that were significantly connected in multiple specific molecular networks (STRING Whole Network p-value=2.2×10^−10^; Fig. 4). Gene-annotated HUI VPFIs were enriched in essential gene ontology (GO) categories or pathways, including: extracellular matrix organization (GO.0030198, p-value=7.0×10^−23^); axonemal dynein complex (GO.0005858, p-value=1.4×10^−11^); proteasome complex (GO.0000502, p-value=2.3×10^−10^) and protein ubiquitination (GO.0016567, p-value=3.9×10^−6^); glycerolipid metabolism (KEGG.00561, p-value=1.9×10^−7^) and lipoprotein transport (GO.0042953, p-value=6.7×10^−5^); 90S preribosome (GO.0030686, p-value=8.1×10^−5^); and cell adhesion molecules (KEGG.04514, p-value=0.00023), which are involved in the response of the immune system to viral pathogens (**Additional file 4: Table S11**).

**Fig. 4.**
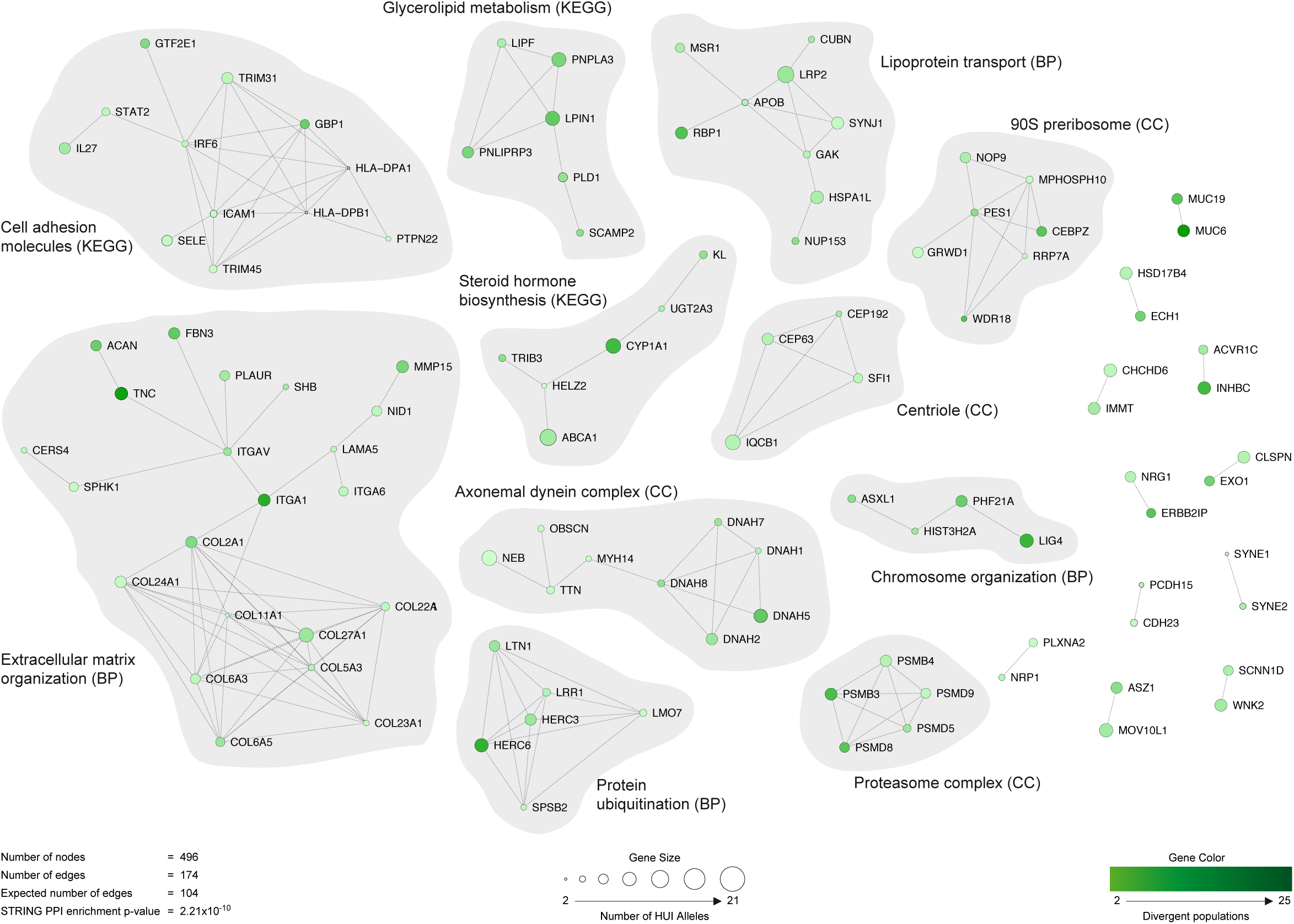
Functional networks between divergent VPFIs detected in Mapuche-Huilliches. Each node in the network represents a gene containing at least 1 highly divergent Fst variant. The scale of green colors (light to dark green) represents the number of 1kGP-phase 3 populations (less to more, respectively) where the variant is divergent. Node size (small to large) indicates the number of alleles present in the sequenced HUI individuals (less to more, respectively). Interactions observed between genes correspond to the information retrieved from the STRING database. Subnetwork names in grey clouds correspond to the most significant ontological/pathway categories obtained from the respective databases.

To detect novel VPFIs we filtered HUI variants through the two-step prioritization protocol described above. Starting with 68,615 novel variants (HUI alleles ≥ 2, no singleton/no monomorphic), 145 surpassed CADD, fell within exonic boundaries and annotated to same number of genes (**Additional file 5: Table S12**). No significant protein-protein connectivity was observed among these products (STRING, highest confidence = 0.9, PPI enrichment p-value=0.303). Nonetheless, novel VPFIs increased the number and further connected the networks of interacting proteins (STRING, highest confidence = 0.9, PPI enrichment p-value=2.2×10^−9^) previously observed with Fst-derived HUI_95th_ variants (**Additional file 1: Fig. S6**). Most notably, the extracellular matrix organization network increased both the number of direct interacting partners and also connected with genes from the glycerolipid metabolism and lipoprotein transport networks. Other enriched networks similarly increased interacting proteins and connections. We finally analyzed overrepresentation of genes linked to diseases or disorders using available information on DisGeNET database [35] and found that divergent and novel variants clustered in networks associated with diseases, most notably in Hypercholesterolemia (umls:C0020443, adj_pvalue=0,00073). Altogether, these findings reveal that HUI genomes contain highly divergent and novel VPFIs that converge in essential ontological categories, which may contribute to common traits, metabolic conditions or prevalent disorders in HUI Native Americans.

## Discussion

We report here for the first time the complete genome sequences of a group of 11 Mapuche-Huilliche individuals and describe common and novel SNVs and large-scale structural variants. Global ancestry composition revealed that HUI genomes have a minimal contribution of European, East and South Asian and African founder populations (K=5) and therefore represent and original source of genetic variation for modern admixed individuals living in America. We found that contemporary American populations, including admixed Latinos from Chile and PEL, have a high component of a previously unknown ancestral genetic contribution identified only in HUI genomes. Such contribution decreased in other Amerindian groups represented in MXL, CLM and PUR populations, likely due to the demographic history of Central and North America. Importantly, in these AMR populations we detected a large genetic component that was not present in HUI genomes and that is also marginally represented in Chilean Latino individuals (gray color in K=10 admixture model), suggesting that AMR ancestors once they reached and settled in the Southern Cone remained isolated and did not mix with other ancestral groups inhabiting the northern tip of South America until very recently, when Spanish conquerors arrived to America. Such idea is further supported by PCA analyses, mtDNA haplogrous, and Fst analyses, which show that HUI individuals are positioned in a narrow cluster at the edge of the distribution of AMR populations.

Native Americans, particularly those from the Southern Cone, are a neglected group in population-based epidemiological studies and remain poorly investigated at the whole genome level [4]. We found highly divergent and novel HUI VPFIs significantly enriched in essential cellular metabolic processes that may help us to understand the differential susceptibility of Mapuche-Huilliches to prevalent metabolic conditions. It has been observed that Native Americans and Latin populations with Amerindian heritage exhibit a substantial predisposition to dyslipidemias, diseases of the circulatory system and neoplasms [14, 36, 37]. We note that the most significant network detected in our study assembled around genes carrying VPFIs in extracellular matrix molecules, glycerolipid metabolism and lipoprotein transport components. Since lipid metabolic pathways tightly control cholesterol homeostasis, it is likely that functional variants altering these cellular processes are involved in triggering lipid-related metabolic disorders in admixed Latin American individuals with Mapuche ancestry.

In summary, here we present the genome sequence of a marginally admixed Native American population from the Southern Cone. HUI genomes show highly divergent large-scale structural and single nucleotide variants that may contribute to modulate susceptibility or resistance to develop complex common diseases in the Chilean and Native American populations. Our study represents an important resource providing a reference panel for Native American populations to be used in future population-based studies on traits of interest (i.e. GWAS with a Native American SNV panel), as well as in the design of early diagnostics and prevention tools.

## Methods

### Whole-genome sequencing

Mapuche-Huilliche individuals from Huapi Island are part of an ongoing longitudinal ultrasonographic study on the prevalence and risk factors of common metabolic diseases in Chile. Mapuche-Huilliche DNA samples were sequenced with the combinatorial probe-anchor ligation sequencing process of Complete Genomics [19]. The standard Complete Genomics bioinformatics pipeline (Assembly Pipeline version 1.10 and CGA Tools 1.4) was used for sequence alignment, read mapping, assembly, and variant call. Human reference genome used was GRCh37.p5 (ftp://ftp-trace.ncbi.nih.gov/1000genomes/ftp/technical/reference/).

### Genotyping of admixed Chilean Latinos

We used a Chilean Latino panel composed of individuals belonging to the ANCORA family health centers located in Santiago-Chile (La Florida and La Pintana) that constitutes an admixed (European-Amerindian) population aged between 20-80 years old from an urban area and representative of the Chilean general population. All subjects were genotyped under the AXIOM^®^ Genome-Wide Platform (version LAT 1) using the GeneTitan^®^ Multi-Channel (MC) Instrument following manufacturer instructions. Samples with discordant sex, elevated missing genotypes rate (≥0.03) or outlying heterozygosity rate (>3SD) were excluded.

### 1000 Genomes Project Phase 3 samples used in our study

The complete dataset from the 1kGP-phase 3 was obtained through the NCBI FTP site (http://www.ncbi.nlm.nih.gov/public/).

### Identification of total SNVs and Indels

Genomic variants were obtained from MasterVar archives delivered by Complete Genomics (MasterVar file description in http://www.completegenomics.com/documents/DataFileFormats_Standard_Pipeline_2.5.pdf). Only high quality reads were used (low_coverage and half variants were filtered). We registered zygosity for every genomic variation: homozygous variations (both alleles are the same and are different from the reference), heterozygous-reference variations (one of the alleles is different from the reference) and heterozygous-alternative variations (both alleles are different and are different from the reference). Variant Call Format files (VCF) were generated from all 11 MasterVar files using the CGAtools software v.1.8 (http://cgatools.sourceforge.net/). Novel SNVs, Insertions and Deletions were defined as genomic variants that are not included in dbSNP build 144 (http://www.ncbi.nlm.nih.gov/SNP) or that have no frequency reported in the 1kGP-phase 3 Database (http://www.1000genomes.org/home) [2], the Exome Sequencing Project (http://evs.gs.washington.edu/EVS/), the Exome Aggregation Consortium (http://exac.broadinstitute.org/), and 46 whole-genome sequences from the Complete Genomics public data (http://www.completegenomics.com/public-data/), which were extracted using the ANNOVAR software [38]. Circular representation of SNVs, CNVs and SVs across HUI genomes was drawn with Circos [39].

### Validation of variants

Variant calling was validated by microarray genotyping using the Illumina Infinium^®^ Human Core Exome BeadChip. The chip consisted in 538,448 variants of which 537,385 are SNVs and 1,063 are indels; 263,929 variants fall in exons. Genotyping study was performed in 9 of the 11 whole-genome sequenced individuals: GS000011194, GS000011195, GS000011196, GS000011198, GS000011200, GS000011201, GS000011215, GS000020403 and GS000020711. The comparison was performed using Variant Call Format (VCF) files which were generated from the Illumina raw genotyping data (Final Report format) taking the genomic positions (chromosome, base pair), the reference allele for the SNV extracted from the NCBI GRCh build 37 reference human genome and the alternative allele from the beadchip annotation data provided by Illumina. The genotype for each SNV (reference homozygote: Hom Ref; reference heterozygote: Het-Ref and alternative homozygote: Hom Alt) was obtained taking the Allele1 and Allele2 Plus information from the raw genotypes. Variants with a Gene Call score (GC) equal or above 0.15 were taken as confident calls, as reported elsewhere [40, 41]. The concordance percentage (Conc%) for each individual was obtained taking the number of matching variants and calculating the percentage according to the total number of matching positions.

### Identity by descent (IBD) and inbreeding analysis

The IBD proportion for each pair of the 11 HUI individuals was calculated in PLINK v.1.9, using a subset of 391,284 SNVs that had > 90% call-rate and were shared between HUI and 1kGP-phase 3 individuals as described before [42]. Linkage disequilibrium (LD) pruning was applied with PLINK so that no pair of SNPs within a 50 SNPs window present an r^2^ value greater than 0.2. Likewise, inbreeding coefficients were calculated in PLINK v.1.9 for each HUI individual using the same subset of LD-pruned SNVs. IBD and Inbreeding analyses shows little cryptic relationships among the samples used in this work. IBD results show no duplicate individuals of first-degree relationships among our sample (IBD > 0.5). Only two couples of individuals (GS000011194-GS000011201 and GS0000111-GS000012210) show second-degree relationships (IBD 0.25, 0.23, respectively). The inbreeding F coefficient analysis shows only 1 individual above the cutoff of −0.12 (GS000012242, F= −0.14). Negative inbreeding values indicate an excess of heterozygosity or “outbreeding” which could happen because of sample contamination, admixture, or genotyping errors. In this case, this particular HUI sample has the highest European (16.5%) and African ancestries (4,4%) among all 11 individuals. These results indicate HUI individuals sequenced are not closely related or inbred.

### Identification of structural variants

Large-scale structural variants (SVs) were identified by two independent methods: The first one is specifically designed to detect copy number variants (CNVs) based on sequence coverage among samples using a Complete Genomics Hidden Markov Model (HMM) that detects significant abnormal coverage over sliding windows. Under the assumption that a sample is diploid, the method can determine if a genomic segment behaves as a “Gain” or “Loss” in comparison to the reference genome (GRCh37.p5) providing Ploidy-number of times the genomic segment in present– and a PHRED-like score which denotes the confidence of the call (computed as −10*log_10_ of the probability of the assigned call being wrong). When whole genome coverage variability is greater than expected, the sample is assigned to a “no-call” state, which impede CNV analysis. The second method uses the CGATOOLS junctions2events pipeline and is based on junctions analysis – defined as regions of the genome where sequences are not adjacent or in the same orientation as present in the reference genome – that rationalize junctions sets in to the following event types: Deletions, Distal Duplications, Tandem Duplications, Inversions and Translocations. Novel CNVs and SVs events were defined as a variable genomic segment present in at least one sample that did not intersect with any region reported in the inclusive map of the Database of Genomic Variants [23] or any of the structural variants reported in the latest release of the 1000 genomes project [24]. Genes affected by structural variants were annotated using Refseq genes from the UCSC table browser and classified into the following categories regarding their overlap context with a structural variant: 5-Prime, 3-Prime, Internal, Complete and Chimeric.

### ADMIXTURE and Principal Component Analysis (PCA)

We selected a subset of 105,252 SNVs with common and eligible genotypes within the 2,504 unrelated samples from the 1000 genomes project phase 3 population and the Chilean panel composed of 1,191 samples giving a final total of 3,706 samples to be analyzed. LD pruning was performed with PLINK so that no pair of SNPs within a 50 SNPs window present an r^2^ value greater than 0.2. This dataset was then introduced to the ADMIXTURE [26] software under default parameters exploring from K = 1 to K = 15 models. Cross validation errors values (CV error) were extracted from each iteration and plotted with R statistical software. Next, this dataset was used to perform a principal component analysis to model ancestry differences between populations using smartpca from EIGENSOFT 5.0 software with default settings [43, 44].

### Maternal ancestry analysis

The complete sequences of mitochondrial DNA of the 11 Mapuche-Huilliches were obtained from the sequencing performed by Complete Genomics. Additionally, 1,016 bp corresponding to the mtDNA control region (rCRS positions 16032-16544 and 051-555) were amplified, purified and Sanger sequenced by Macrogen, South Korea as described [15, 29]. Sequences were aligned and edited with Alignment Explorer (MEGA 4.0)[45]. There was complete concordance between Sanger sequencing and mtDNA genome sequences provided by Complete Genomics. Polymorphisms were confirmed directly using Sequencher 4.9 vDemo (http://genecodes.com/). Sequences were grouped by mitochondrial haplogroup and analyses were performed separately. The results were confirmed by comparison with mtDNA tree Build 15 (rCRS-oriented version of Build 15) available on the PhyloTree.org website. Mitochondrial DNA haplogroups from different Native-Chilean and other southern South American populations have been described elsewhere [15, 29]. Calculations were performed using the Network 4.5.0 program (http://www.fluxusengineering.com/sharenet_rn.htm); median joining and maximum parsimony were used as post-processing options.

### Fixation index (Fst) analysis in HUI genomes

Fst analysis was performed using the Weir & Cockerham's Fst estimator (wcFst) [31] function inside VCFTools software V.0.1.12b [46]. We obtained Fst statistics at population (weighted Fst) and at SNV level from the comparison between HUI and all individuals from each of the 26 populations of the 1kGP-phase 3: Africa (AFR: YRI, LWK, GWD, MSL, ESN, ASW and ACB), Europe (EUR: CEU, IBS, GBR, FIN and TSI), America (AMR: CLM, PUR, MXL, and PEL), Southern Asia (SAS: PJL, GIH, BEB, STU and ITU) and East Asia (EAS: CHB, JPT, CHS, CDX and KHV). Briefly, to get genotype homogeneity between all HUI individuals, we merged no singleton and no monomorphic SNVs with call rate above 90% with the whole panel of genotypes from 1kGP-phase 3 (also filtered using Vcftools V.0.1.12b) to get a set of common variants between all populations. In this subset of variants, we calculated the number of divergent SNVs per population using the genome-wide Fst distribution and setting a 95th percentile cutoff as a divergence threshold. Violin plots for the population level Fst distributions were done using the ggplot2 R software [47, 48].

### Identification of VPFIs with CADD, STRING networks

Genetic variants were queried for functional impact using ANNOVAR [32] with the Combined Annotation Dependent Depletion (CADD) database [33]. We selected variants surpassing the Phred-Scale score of 20, which represent the top 1% of most deleterious substitutions, and defined them as Variants with Potential Functional Impact (VPFI). Protein-protein interaction connectivity among products encoded by divergent and novel HUI VPFIs was examined using the online database resource Search Tool for the Retrieval of Interacting Genes (STRING) [34]. Significant networks were visualized using Cytoscape 3.5.1 (http://www.cytoscape.org).

## Declarations

### Ethics approval and consent to participate

Mapuche-Huilliche individuals from Huapi Island are part of an ongoing longitudinal ultrasonographic study on the prevalence and risk factors of common metabolic diseases in Chile. Informed consent for the study of genetic and metabolic risk factors for prevalent metabolic diseases was obtained from all studied participants in years 1993 and/or 2001 [49, 50]. Oral and written informed consent from the legal representatives of Huapi Island Mapuche-Huilliche community for whole genome sequencing in some members of the community was obtained in January 2012. The present study was approved by the Institutional review Board for Human Studies of the Faculty of Medicine at Pontificia Universidad Católica de Chile.

### Availability of data and material

Whole genome sequencing data used in this manuscript is available at the Sequence Read Archive (SRA, https://www.ncbi.nlm.nih.gov/sra), under accession number SUB2180735, BioProject ID at http://www.ncbi.nlm.nih.gov/bioproject/358028.

## Competing interests

The authors declare no conflict of interest.

## Funding

This work was funded by grants from Fondo de Areas Prioritarias (FONDAP) Center for Genome Regulation (number 1509000), FONDAP Center for intercultural and indigenous research (number 15110006), Fondo Nacional de Desarrollo Científico y Tecnológico (FONDECYT) numbers 1130303 to J.F.M., 1140353 to G.V.D. and 1120813 and 1160833 to S.E.

## Authors’ contributions

RAG, JFM, and GVD designed the study; EAV, TCM, BIB, EP-P, CM, AM, LA, DS, ER, MV, AD and MM developed analytical tools, performed analysis and interpretation of the data. KP, PN, SB, JH, CH, CM, AR-J, DT, FV, PV, MM, SE, analyzed the data. MLA, VC, MG, MM, AO and AM helped in drafting the manuscript. EAV, RAG, JFM and GVD wrote the paper.

## Acknowledgements

The authors thank the Mapuche-Huilliche community of Huapi Island (Ranco Lake), without whose participation and encouragement this work would not have been made possible.

